# The wheat head blight pathogen *Fusarium graminearum* recruits facultative endohyphal bacteria from the soil, making the fungal-bacterial holobiont nitrogen-fixing and increasing the fungal pathogenicity

**DOI:** 10.1101/2022.06.24.497425

**Authors:** Hina Ali, Mengtian Pei, Hongchen Li, Wenqin Fang, Hamid Ali Khan, Tariq Nadeem, Stefan Olsson

## Abstract

In nature, fungal endophytes often have facultative endohyphal bacteria. Can a fungal pathogen such as *Fusarium graminearum*, pathogenic on wheat, get facultative endohyphal bacteria from soil (FEB) and how does the FEB affect the fungal phenotype? We constructed a growth system/microcosm that allowed a molecularly well-studied *F. graminearum* isolate, PH-1, to grow through natural soil and then be re-isolated on a gentamicin-containing medium, allowing endohyphal growth of the bacteria while killing eventual bacteria growing on the agar medium. We had labelled the *F. graminearum* PH-1 with a *His1mCherry* gene staining the fungal nuclei fluorescent red to confirm re-isolation of the same isolate we sent through the soil. Through qPCR of the *16SrRNA* gene in the bacteria using universal primers combined with qPCR of the *mCherry* gene of DNA from the re-isolated cultures of the Fg-FEB holobionts growing on gentamicin-containing media, it was found that most of the holobiont isolates contained about 10 *16SrRNA* genes per fungal *mCherry* gene. The Fg-FEB holobiont isolates were sub-cultured several times, and the FEB content on lab media was stable. Sequencing the *16SrRNA* gene from several Fg-FEB holobiont isolates revealed known endophytic bacteria capable of nitrogen fixation. We compared the pathogenicity of one of the Fg-FEB holobionts *Fg-S.maltophilia*, with the background without FEB and found that it was more pathogenic than without FEB. We could also show that the bacterial *16SrRNA* load per fungal *His1mCherry* gene inside the wheat stayed the same as in culture. Finally, we tested if the *Fg-S.maltophilia* was capable of nitrogen fixation and could show that it, on a nitrogen-free medium, formed a dense mycelium containing proteins at similar levels as on regular nitrogen-containing media. Our results could indicate that naturally occurring fungal pathogens outside lab conditions might contain facultative endohyphal bacteria, positively affecting their pathogenicity and ecological fitness.

## 1. Introduction

Fungal-bacterial interactions are old and can be different from simple competition for space and nutrients to more intimate interactions when bacteria grow on the surface of fungi, more or less dependent on nutrients. Bacteria can also predate or parasitize fungi or live inside the fungi as obligate or facultative symbionts (Frey-Klett et al., 2011; Olsson et al., 2017; Deveau et al., 2018; Junier et al., 2021)

Most fungi in nature might have facultative endohyphal bacteria (FEBs). These Endohyphal bacteria (EHBs) are widespread in rhizosphere fungal phyla like Basidiomycota, Glomeromycota, and Mucoromycotina, especially in the highly diverse Ascomycota, endophytes of stem, root, and leaves, class 3 endophytes (Rodriguez et al., 2009). EHB can influence the phenotypes of root-associated fungi and shape the outcomes of plant-fungal interactions (Ghignone et al., 2012). EHBs are host specific, vertically, or horizontally transmitted and frequently maintain obligate relationships with fungi in the rhizosphere (Mondo et al., 2012). In contrast, associations between EHB and many species of Ascomycota seem to be facultative (FEB) (Hoffman et al., 2013). However, many plant-associated microbes remain undescribed (Li et al., 2014) and are found in the association between with fungi as endosymbionts (endobacteria, endofungal bacteria, or Endohyphal bacteria (EHB) (Márquez et al., 2007; Partida-Martinez et al., 2007; Naito et al., 2015). Besides, these EHBs can be re-isolated from their hosts by antibiotic treatment and are often cultured on standard selective nutrient media (Hoffman et al., 2013) and should be labelled as FEBs. However, functional relationships have been studied for only a few associations, limiting inferences regarding the scope and potential importance of EHB-fungal associations in ecological interactions and human applications (Frey-Klett et al., 2011; Lackner et al., 2011).

Many bacteria can move by swimming, swarming, twitching, sliding, or gliding without pili. These bacteria form a synergetic film around the fungal hyphae and disperse along with fungal mycelium growth in natural environments such as soil which act as fungal highways (Kohlmeier et al., 2005). This mycelium-driven transport plays a vital role in regulating the distribution of bacteria in unsaturated soils. In vitro studies have also emphasized the ecological benefits of this dispersal mechanism. For example, to enhance the biodegradation of soil pollutants, maintenance of bacterial flagella and long-term storage of carbon and soil fertility by fungal-bacterial interactions in the context of the oxalate-carbonate pathway has been demonstrated in soil microcosms (Simon et al., 2015). However, evidence of active bacterial movement along mycelium in soils is still missing despite its potential ecological importance. Various approaches have been used to isolate selectively soil bacteria moving along fungal mycelium. One is a microcosm model based on a labelled Petri dish, pre-inoculated with Lyophyllum sp. strain Karsten to identify numerous bacterial species potentially driven along the fungal hyphae (Simon, 2016). More recently, an inverted Petri dish method in which *Pythium ultimum* (an Oomycete and not a fungus) mycelium was inoculated in the soil as a “fungal highway” to isolate contaminant-degrading bacteria (Furuno et al., 2012). Subsequently, another approach was developed to isolate bacteria from the hyphae of filamentous microorganisms using fungal highway columns without pre-inoculation (Junier et al., 2021).

This study aimed to re-introduce Facultative Endohyphal Bacteria (FEBs) from the soil into *Fusarium graminiarum* PH-1 hyphae and the effect FEBs on the pathogenicity of the fungal-bacterial holobiont (Deveau et al., 2018). A reporter strain was constructed by transforming the *mCherry* gene (red fluorescent) fused with the Histone1 linker inserted into the PH-1 WT strain. The reporter strain (FgHis1mCherry) was sent through the soil using a slightly modified fungal highway column setup (Junier et al., 2021) and re-isolated on a selective medium (DFM medium containing gentamicin (0.52mg/l) along with the potential FEBs. Several FEBs were found, and one potentially nitrogen-fixing FEB fungal bacterial holobiont, Fg-*S. maltophilia* (Table S1 explains how we name fungal holobionts and constructs in this work) was investigated in more detail. It contained the FEB in both mycelia and conidia inside a plant during infection, increased pathogenicity, and fixed nitrogen. The possibility that most fungal pathogens contain EHBs in nature is discussed.

## 2. Methodology

### 2.1 Fungal strains, culture, and growth conditions

The *F. graminearum* strain PH-1 grown on DFM agar medium (Frandsen et al., 2010) was used as the wild-type strain. PH-1 and the FgHis1mCherry reporter strain were used as a non-bacterial background strain, and the Fg-FEB fungal strains containing FEBs were grown and stored on DFM solid medium (Malz et al., 2005) supplemented with Gentamicin (512 μg/mL). The fungal cultures were incubated at 24°C with 12 hours light and dark cycles. For long-term storage, the fungal cultures on small cut pieces of Whatman cellulose filter paper were stored in sterilized envelopes on silica gel at 4 °C. **Table S1** contains descriptions and explanations for naming gene constructs and strains.

**Table S1.** The naming of constructs and strains, including names for fungal-bacterial holobionts. Fg= *Fusarium graminearum*

| Description | Abbreviation (in figures) | Explanation |
| --- | --- | --- |
| Fg containing FEBs | Fg-FEB | Short for the fungal species and FEBs of any bacteria. FEB since the endohyphal bacteria were acquired from the environment |
| Fg containing the histone 1 gene fused with the mCherry gene so that it can be recognized as the strain we re-isolate. | *FgHIS1mCHERRY* | When referring to having the gene construct or the gene expression. Gene in italic. The gene is however the orthologous gene *MoHIS1mCHERRY* pointed out in methods |
| Fg strain containing the histone 1 gene fused with the mCherry gene when discussing protein | FgHis1mCherry (FgmCherry) | When referring to the reporter protein carrying strain. The protein is however a MoHis1mCherry. |
| Plasmids | *pCB1532* | Referring to the plasmid. DNA so gene in italic. |
| Fg containing:<br>*Stenotrophomonas pavanii*<br>*Stenotrophomonas maltophilia*<br>*Phytobacter urisingii* | Fg-*S.pavanii*. (Fg-Sp)<br>Fg-*S.maltophilia* (Fg-Sm)<br>Fg-*P.urisingi* or (Fg-Pu) | The Fg isolates that are known to contain the specific isolates. |
| 16SrRNA gene<br>MomCHERRY gene | *16SrRNA*<br>*mCherry* | For qPCR |

### 2.2 Construction of reporter strain and validation of (FgHis1mCherry) reporter strain

A fungal reporter strain (FgHis1mCherry) was constructed by labelling the *F. graminearum* PH-1 with the vector *HIS1-mCHERRY-pCB1532* previously constructed in our lab for the transformation of *Magnaporthe oryzae* (Zhang et al., 2019). The linker protein histone His1 in the relatively closely related to *F. graminearum* is very similar. Positive fungal transformants were selected on DFM agar medium. The FgHis1mCherry reporter strain was validated for the presence of the vector in one copy. The gDNA isolated from the FgHis1mCherry reporter strain was amplified using PCR with the help of specific primers, designed by primer5 software according to the following the PCR steps: Initial denaturation for 5 min at 95 °C, Denaturation for15 sec at 95 °C, Annealing for 15 sec at 58 °C, extension for 1 min and 30 sec at 72 °C and Final extension for 10 min at 72 °C. The PCR product was run in 1% Agarose Gel to check the visible bands for the FgmCherry reporter strain. The eventual change in morphology phenotype of the FgHis1mCherry reporter strain compared to PH-1 was evaluated on DFM solid medium plates and kept at 28°C in an incubator for 5 days. The expression of FgmCherry protein in the reporter strain was validated through confocal microscopy for the presence of red fluorescent nuclei using a NikonA1 confocal microscope and 100X magnification (Plan Apo VC 100x Oil DIC,NA=1.40), and it was also used to confirm that the hyphae are typically nucleated. Most importantly, the FgmCherry reporter strain was tested for conidiation, sexual reproduction, and pathogenicity assay and compared with PH-1 to test that the random integration of the *HIS1-mCHERRY* construct has not disrupted any genes needed for wheat infection.

### 2.3 Development of an isolation method to catch FEBs using a Fungal highway column technique

A fungal highway column with modifications to isolate FEBs was developed based on an already described technique for selectively isolating bacteria moving following the fungal hyphae (Simon et al., 2015), combined with using Gentamicin to keep the FEB inside the hypha (Baltrus et al., 2018). The fungal highway column was designed as a closed device (Ø: 15 mm, height: 48 mm) intended to isolate bacteria able to grow along or inside the fungal mycelium towards a target culture medium containing DFM + Gentamicin killing bacteria on the hyphal surface but leaving intracellular bacteria unharmed. The test tube and lid were sterilized by immersion in 70% ethanol for 30 minutes and exposed to UV-C light for 30 minutes before assembling the system. Culture media and glass beads were sterilized by autoclaving (21 min at 121°C). Each column was assembled inside a sterile laminar flow hood. At the bottom of the fungal highway column, the first layer of water-agar was added to fill the rounded bottom of the column to make flat support and was allowed to solidify. Then a second thin layer of melted DFM medium was poured on the top of the water agar and allowed to solidify and cool. A small mycelium plug of the FgHis1mCherry reporter strain from the DFM agar plate was taken with the help of a sterilized surgical blade and inoculated onto the centre of the surface of the DFM agar medium. Next, a thin layer of sterilized glass beads was added on top of the DFM agar disk with fungus to avoid direct contact with the soil that was added next. Then a thin layer of sieved moist soil was added, followed by sterilized glass beads on the top of the soil in a thick layer. A target medium, a DFM agar medium disk containing 512 μg/mL gentamicin cut with a sterile cork bore and added on the top of the thick layer of grass beads. The ready column was closed with the sterile cap. The column was incubated at 25°C with 12 hours of light and dark photoperiod. It was inspected regularly and allowed to incubate until the fungus had grown from the inoculum, through the soil and the thick glass bead layer, and onto the target medium, and the fungal growth was visible on the target medium.

### 2.5 Isolation and purification of Fg-FEBs

The facultative endohyphal bacteria (FEBs) moved along with the FgmCherry reporter mycelium are now potential FgmCherry strains containing FEBs (Fg-FEBs) found on the top of target DFM agar medium containing 512 μg/mL Gentamicin. The potential Fg-FEBs were isolated using a Pasteur pipette tip using a slight suction technique. The DFM target agar disc, covered with fungus, was lifted out and inoculated on fresh agar medium with Gentamicin (512 μg/mL) in 10 cm diameter Petri dishes by placing the disc against the agar and releasing the suction. The plates were Incubated until the fungus had grown over most of the DFM agar medium. For purification and to be more sure only to get FEBs, small plugs from the edges of potential Fg-FEB colonies were taken and subcultured on DFM with Gentamicin (512 μg/mL) agar plates. These plates were incubated at 25 °C for 3-5 days with a 12 hours of light and dark photoperiod. After incubation, when plates were covered with the growth, the potential Fg-FEB culture was tested for the presence of FEBs.

### 2.6 Molecular analysis of the presence of FEBs inside the potential Fg-FEB cultures using 16s rRNA gene sequencing

*16SrRNA* gene sequencing PCR used the universal primers for *16SrRNA* gene sequencing and qPCR after confirming that these primers perform quantitatively in a qPCR (data not shown). The primers for qPCR of the mCherry gene were designed with the help of primer5 software (Table 1). The conditions for PCR were: Initial denaturation for 5 min at 95 °C, denaturation for 30 sec at 95 °C, annealing for 30 sec at 58 °C, extension for 1 min and 30 sec at 72 °C and final extension for 7 min at 72 °C. The PCR product was run in a 1% agarose gel to check the visible bands for the presence of FEBs. After gel electrophoresis, the positive DNA samples were purified from the gel using a QIAGEN’s gel extraction kit and sent to the QIAGEN, Fuzhou, Fujian, China for *16SrRNA* gene sequencing to further identify FEBs. After the 16SrRNA gene sequencing, the sequences were used for blast search for the specific identification of FEBs.

**Table 1.**
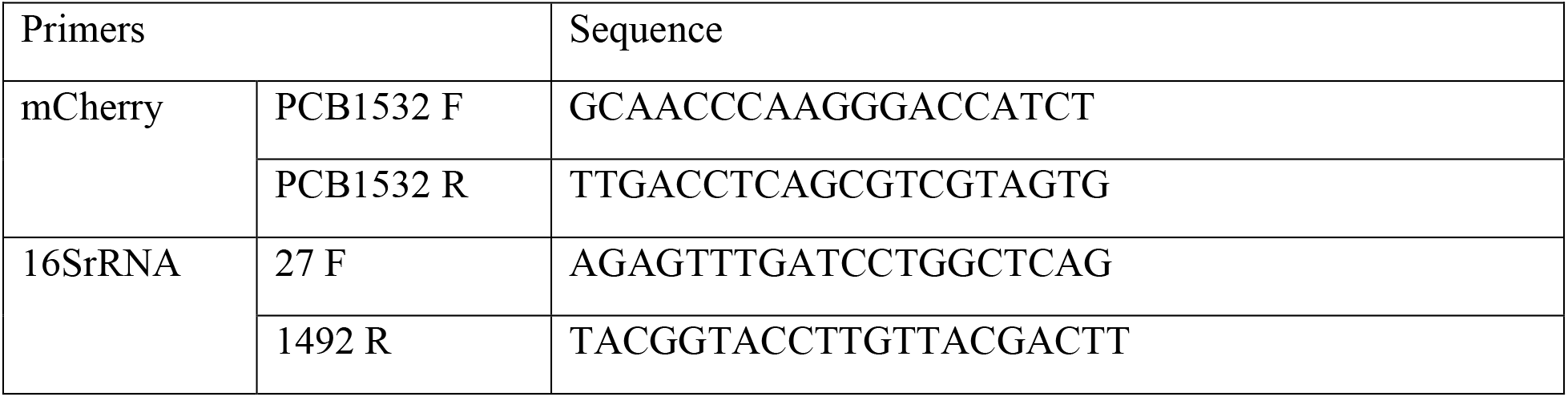
List of Primers for mCherry and 16SrRNA qPCR and 16SrRNA sequencing

### 2.7 Comparison of Fg-FEB colony phenotypes with FgHismCherry

The colony morphology phenotype of the Fg-FEBs cultures was compared with background strain FgHismCherry cultures after growth on DFM agar medium with Gentamicin (512 μg/mL) and incubated at 25 °C for 4-5 days.

### 2.8 qPCR of *FgFEBrRNA/His1mCHERRY* to estimate bacterial load in the fungus

The estimation of FEBs per Fg nuclei in the Fg-FEB strains was done using a qPCR procedure: Initial denaturation at 95 °C for 2 min following 40 cycles, denaturation at 95 °C for15 sec, annealing at 55°C for 15 sec and extension at 60°C for15 sec. The amount of *His1mCHERRY* reporter gene was used as a reference gene, and the ratio of *16SrRNA* gene copies per *His1mCHERRY* reporter gene is an expression of the FEB load per nuclei of the fungus. For qPCR SYBR premix, Ex Taq II system (TaKaRa Perfect Real Time) was used. The DNA isolated from respective Fg-FEB isolates was used as templates.

### 2.9 Pathogenicity assay of *Fg-S.maltophilia* and the usual WT PH-1 used in labs and estimation of fungal bacterial load during wheat coleoptile infection

Wheat coleoptiles were infected with 10 μL conidia suspension, 10 μL conidia suspension of *Fg-S.maltophilia* or PH-1, or using mycelial discs and incubated at 25°C, with 12 hours of light and dark cycles photoperiods, to determine the severity of coleoptile infection of the fungal isolates. The lesion size on infected wheat seedlings’ coleoptiles was measured at 7 dpi by photographing them with a ruler as a reference (Jia et al., 2017). qPCR was employed to estimate the bacterial load as *16SRNA* gene copies per *mCherry* gene copies during wheat coleoptiles infection when conidia and mycelia were used as inoculum. The bacterial load in the Fg-*S.maltophilia* grown in culture on DFM+Gentamicin was used for comparison. For qPCR SYBR premix Ex Taq II system (TaKaRa Perfect Real Time) was used. DNA isolated from the infected wheat coleoptiles was used as a template.

### 2.10 Testing nitrogen-fixing capability of *Fg-S.maltophilia* v/s PH-1 and testing the nitrogen-fixing ability by comparing protein content of cultures

A synthetic agar medium DFM with double glucose content as only added carbon source and no added nitrogen supplemented with Gentamicin at a 512 μg/mL was used to keep the *S. maltophilia* FEB inside the fungus. The plates were incubated at 25 for 15-20 days with a 12 hours of light and dark photoperiod. The Bradford protein assay was used to determine the protein amount of *Fg-S.maltophilia* and PH-1 cultures on the medium without added nirogen source. Protein was extracted from a standardized area of the mycelium using a cork plug, and total proteins were extracted. The amount of Coomassie 1 Brilliant Blue G-250 dye colouring of the extracted proteins (Bradford 1976) was then used to measure total protein content. A dilution series of bovine serum albumin (BSA) 2 mg/mL solution was used as a standard reference. The procedure followed the Quick Start Bradford Protein Assay Kit 1 (Bio-Rad) protocol.

#### Statistics and statistical considerations

In this work, we often use ratios between two parameters measured for the same sample since that is a more stable comparison of these measured parameters and does not depend on random variations in sample biomass due to sampling errors. Since ratios of normally distributed measurements of parameters in the same samples are Log-normal distributed, we chose to calculate the confidence intervals for the log-normal distribution but present the bar plots with these ratios having a log2 y-axis labelled with the actual ratios. That way, the error bars are shown correctly and symmetric around the averages, irrespective of the ratio values. As a final service to the reader, we use 95% confidence interval error bars instead of pairwise t-tests or t-tests to a control. That means that the readers can choose to compare whatever they find of interest, and we are also sure of not overinterpreting and assigning too low a probability for the null hypothesis of no difference in any comparisons. Only averages with non-overlapping 95% error bars are considered where P for the null hypothesis of no difference ≪0.05.

## 3. Results

### 3. 1 Validation of FgHis1mCherry Reporter strain

A reporter strain (FgHis1mCherry was constructed and validated using PCR, colony morphology, conidiation, sexual reproduction, and pathogenicity assay and compared with PH-1. The colony morphology, sexual reproduction, and pathogenicity assay of the FgHis1mCherry strain were similar to PH-1 (**Fig 1 A,B,F-H**). Further, validation of FgHis1mCherry was done by confocal microscopy. Red fluorescence fungal nuclei (in the figure showed using a pseudocolor gradient to highlight the accumulation in the nuclei) showed the presence of plasmid *pCB1532* containing and expressing His1mCherry (**Fig 1 C-E**).

**Figure 1.**
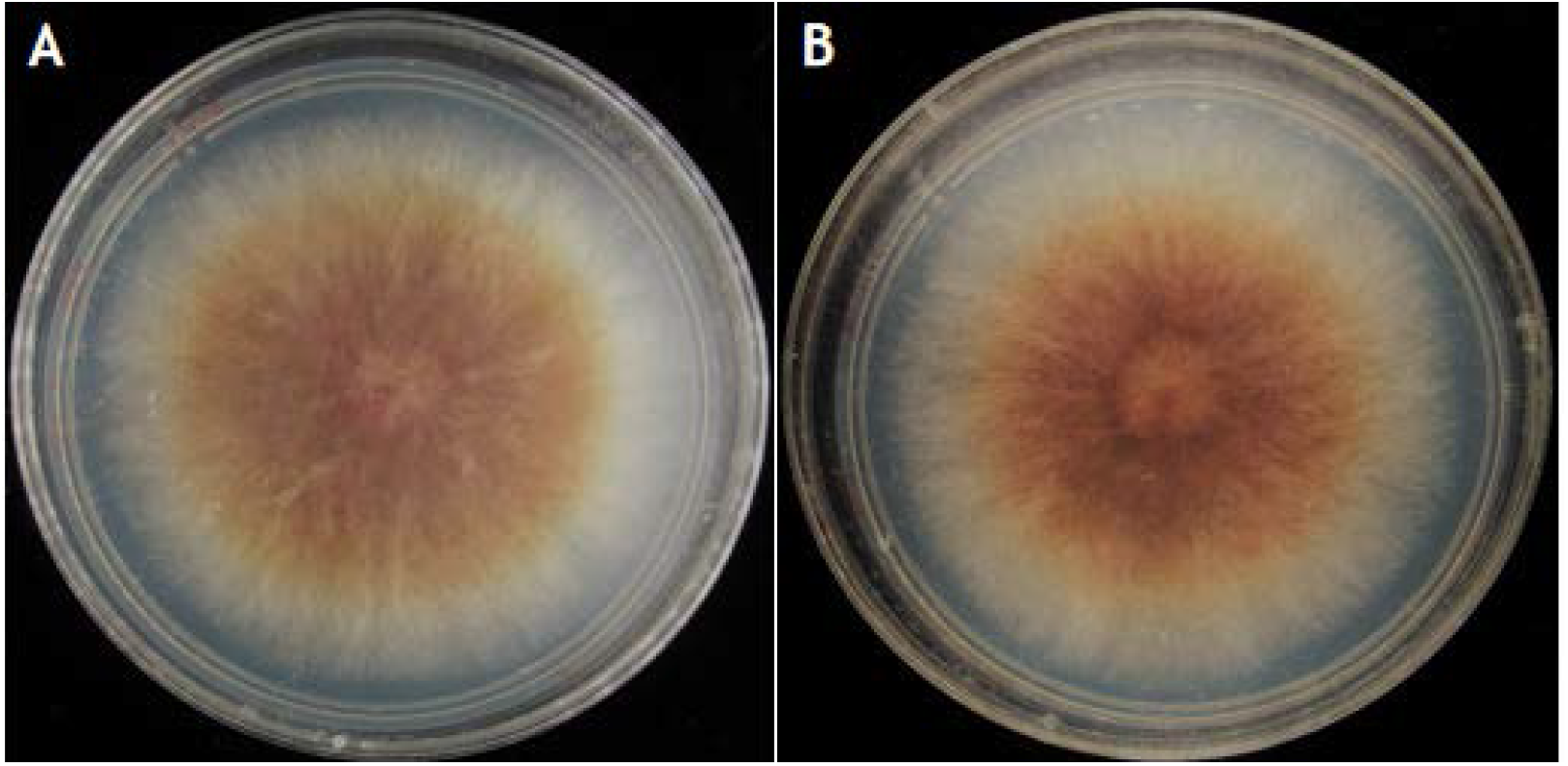
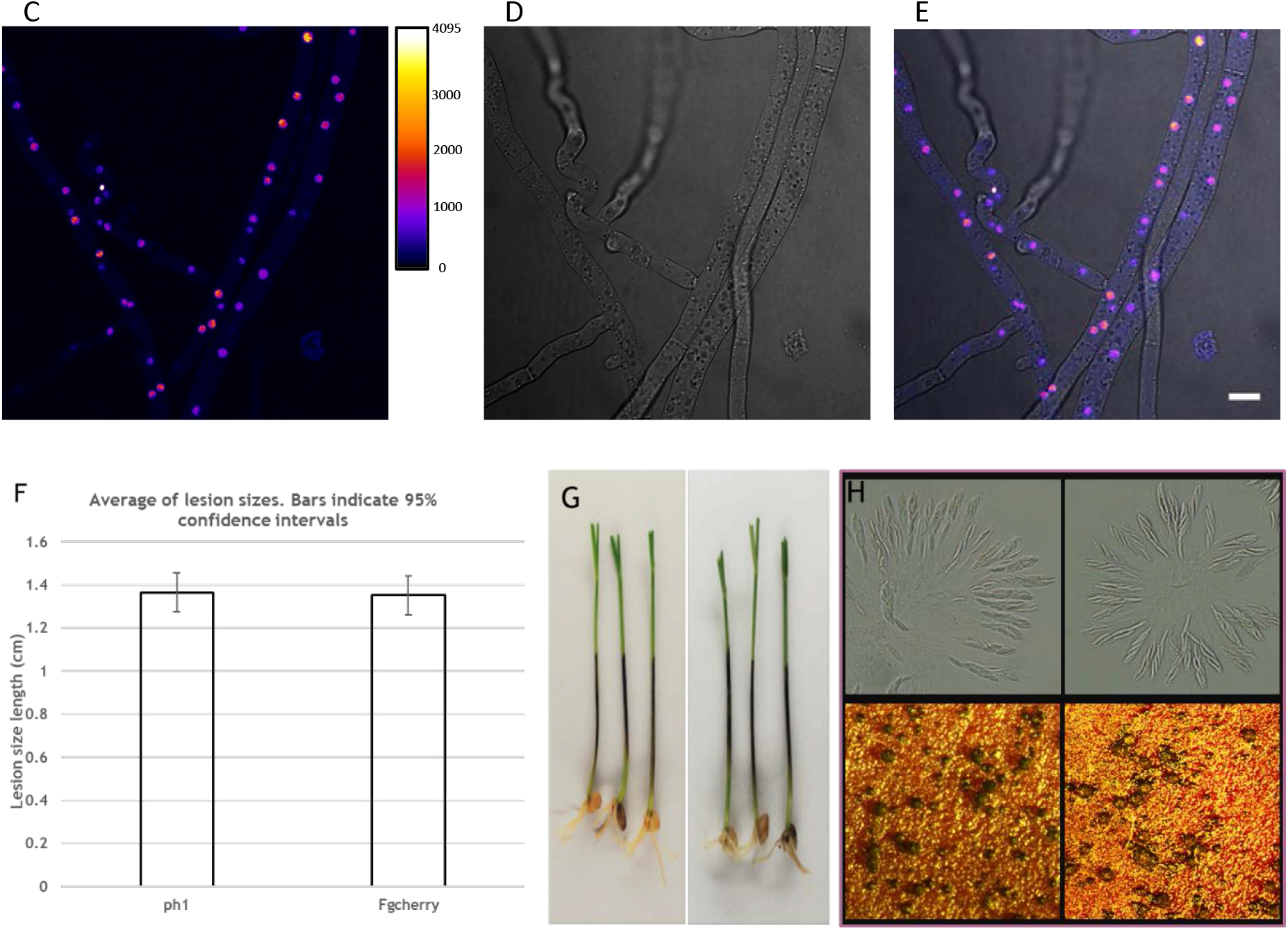
Validation of reporter strain FgHis1mCherry. (A) Wild type PH-1 grown on DFM medium supplemented with gentamicin 512 μg/mL to kill extracellular bacteria and is used for finding intracellular bacteria later. (B) Compared with reporter strain FgHis1mCherry (FgmCherry). (C)(D) Validation of Reporter strain (FgHis1mCherry) shows the presence of the gene product of *HIS1mCHERRY* (*pCB1532*) under a confocal microscope. (E) The red fluorescence *Fusarium graminearum* is shown here using a Fire lookup table to show a large range of intensities of the confocal microscope (4096 levels) as temperature colours range black-blue-red-orange-white to show a large intensity range in a way better adapted to the human eye, showing that His1mCherry mainly accumulates in nuclei. (F) Comparison of lesion size length on wheat coleoptiles between Ph-1 and FgmCherry. The error bar shows a 95% confidence level indicating that bars with overlapping error bars are not likely to be significantly different from the null hypothesis that they are the same. (G) Wheat coleoptile lesions PH-1 (left panel) and FgHis1mCherry (right panel). (H) Ascospore formation is normal for both PH-1 (top left panel) and for FgHis1mCherry (top right panel), as well as perithecia formation for PH-1 (bottom left panel) and FgHis1mCherry (bottom right panel) (DIC image overlay with 10 μm size bar).

### 3.2 Isolation and Purification of Fg-FEB holobionts

A modification of the fungal highway column (Simon et al., 2015) was constructed to obtain Fg-FEB holobionts from soils for the isolation of FEBs coming along with fungal hyphae towards a target DFM-medium supplemented with Gentamicin to isolate (512 μg/mL) (**Fig 4**) to isolate FEBs (Baltrus et al., 2018). After isolation on gentamicin-containing DFM plates, the FgHis1mCherry reporter strain still harboured FEBs when the culture was fully grown on the DFM+gentamicin medium surface. The potential Fg-FEBs were further sub-cultured for purification. The original FgHis1mCherry reporter strain can be reliable prescreened before PCR-final confirmation (below) by using any system using green light illumination and recording red fluorescence since the white colony edge becomes red fluorescent (we used our eyes since the colonies edges become slightly pink to the eye in white light).

**Figure 4.**
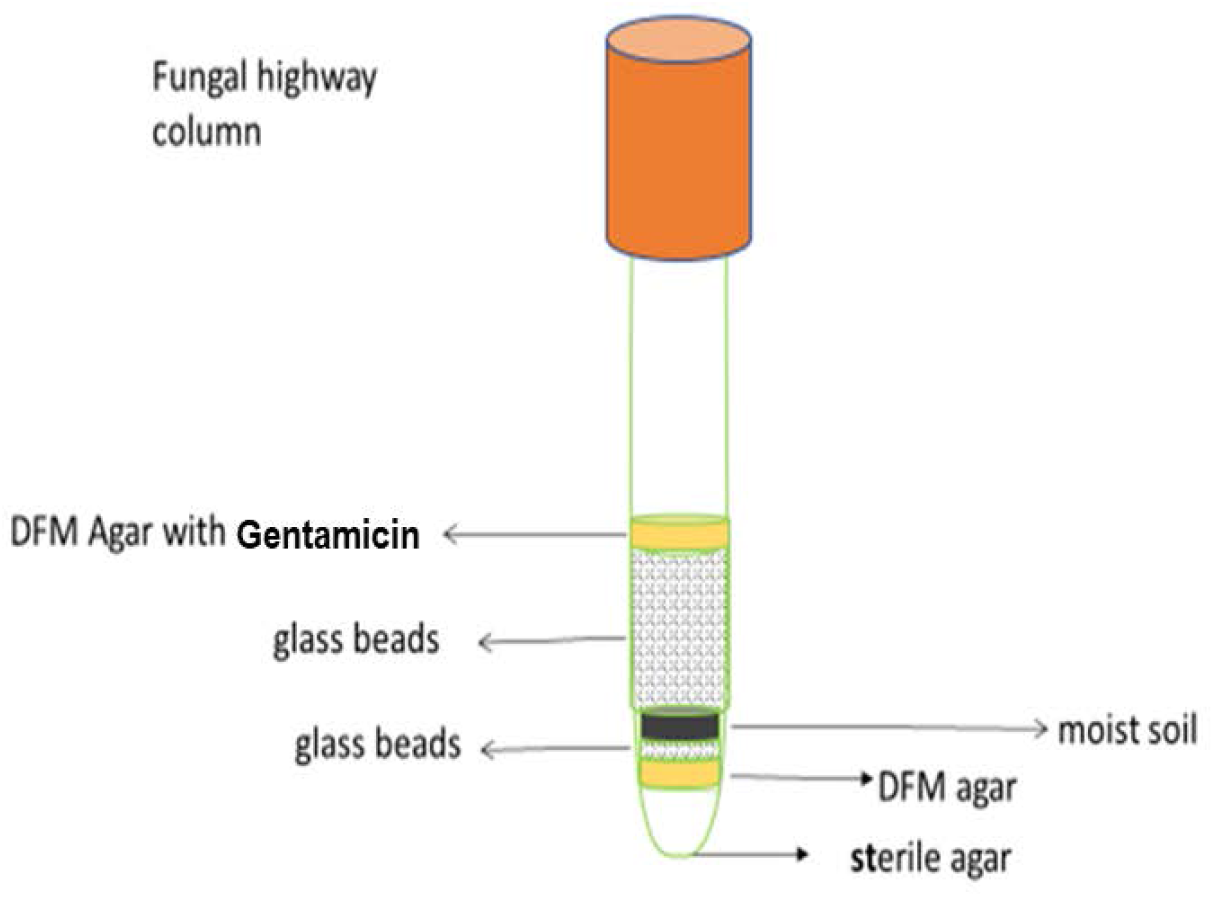
The developed method is based on the Fungal Highway column technique to isolate fungal-associated facultative endohyphal bacteria harboured by *Fusarium graminearum* but should work with many other fungi.

### 3.3 Molecular detection of FEB inside FgHis1mCherry reporter strain

The presence and identity of FEB inside the FgHis1mCherry reporter strain were confirmed using molecular analysis. FEBs inside FgHis1mCherry were detected using PCR of *16SrRNA* genes using the 16S universal primers for bacteria (Table 1). After detecting of FEBs inside FgHis1mCherry, the positive isolates were sent for DNA sequencing. After sequencing, the sequences were BLAST searched on the NCBI 16S sequence database. We found identities with several types of facultative endohyphal bacteria inside the FgHis1mCherry reporter strain. Some commonly occurring were *Stenotrophomonas pavanii (Fg-S.pavanii), Stenotrophomonas maltophilia (Fg-S.maltophilia*, or Fg-Sm for short) and *Phytobacter ursingii Fg-P.ursingii*). These are known endophytes of plants capable and interestingly capable of nitrogen fixation (Ramos et al., 2011; Pillonetto et al., 2018; Alexander et al., 2019). It is not an exhaustive list of bacterial species that can be facultative endohyphal *F.graminearum*, but we continued working with Fg-*S.maltophilia*.

### 3.4 Effect of FEB on FgHis1mCherry colony morphology on standard media

*Fg-S.maltophilia* morphology was compared with the background FgHis1mCherry strain to check the FEB’s effect on the phenotype. The isolates were tested to evaluate the growth compared to FgHis1mCherry. The result indicates that *Fg-S.maltophilia* have a similar colony phenotype as FgHis1mCherry and the FEB has no marked effect on the colony phenotype (**Fig 5**).

**Figure 5.**
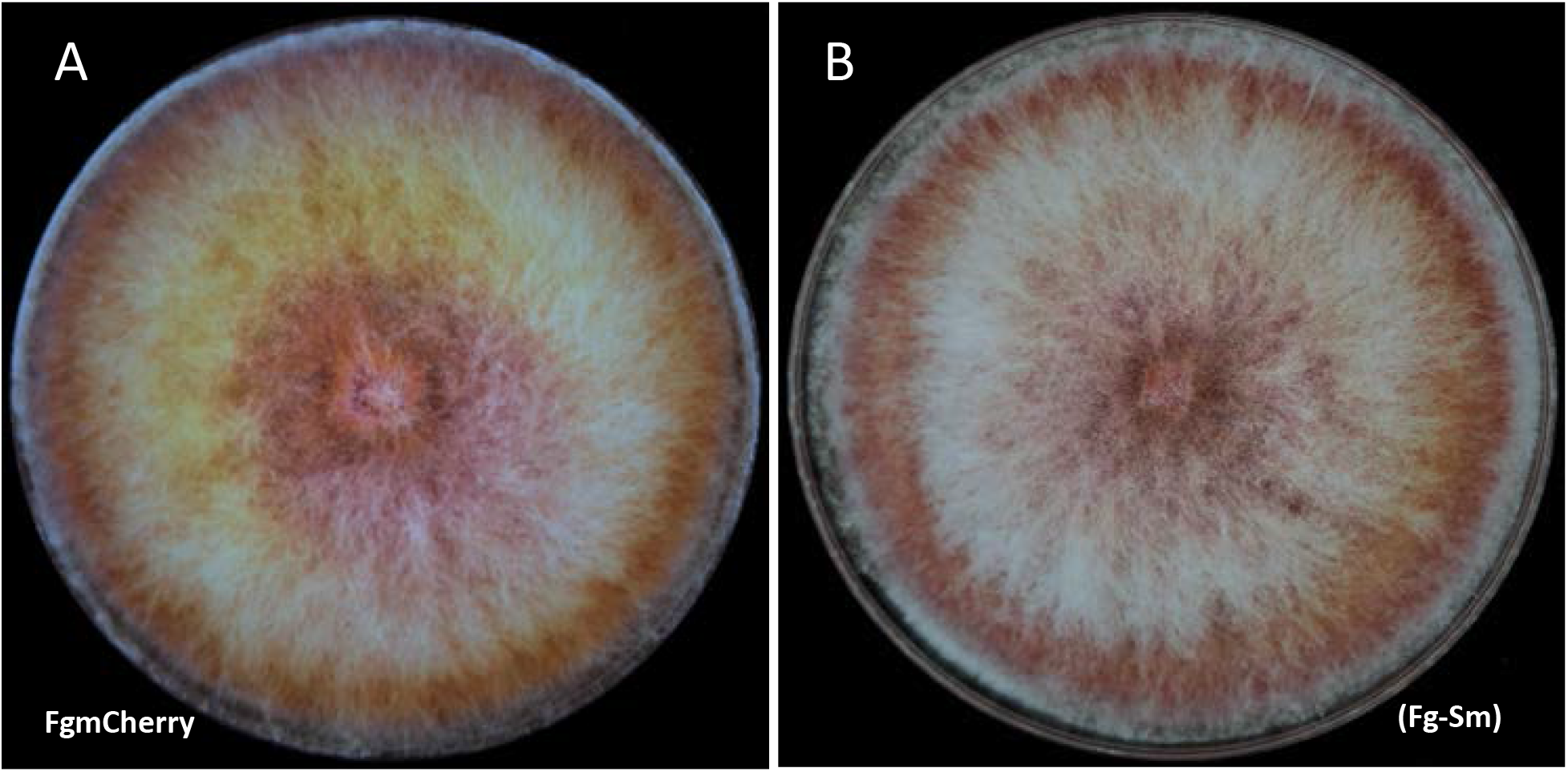
Morphology of FgHis1mCherry strain (FgmCherry). (A) (FgmCherry) compared with the Fg-*S.maltophilia* (Fg-Sm) (B) isolate. Both were grown on DFM medium supplemented with gentamicin 512 μg/mL. Both grew at the same rate and had similar morphology.

### 3. 5 Stability of *Fg-S.maltophilia* and estimation of bacteria load of the fungus

After three months of subculturing (3 times) on a DFM medium supplemented with Gentamicin, the bacterial load was estimated as the number *16S rRNA* genes per *mCherry* genes using qPCR. The subculturing was done in 3 separate lines of transfers between plates to estimate eventual difference over time in the load of bacteria in the fungaus. The load stayed relatively constant, and we found it to be 10^1.18 (95%conf interval 0.567)^ =15 (11-41 95% confidence interval). When growing the Fg-*S. maltophilia* on DFM medium without gentamicin bacterial colonies could be seen among the hyphae, while this was not the case for the sparce growth on distilled water agar.

### 3.6 Pathogenicity of an *Fg-S.maltophilia* v/s PH-1

A pathogenicity assay of Fg-FEBs was conducted and compared with the PH-1. The wheat coleoptiles were infected with conidial suspension or mycelium disc of *Fg-S.maltophilia* and the wild-type PH-1. After incubation at 25°C, lesion size on coleoptiles of infected wheat seedlings was measured at 7 dpi by photographing them with a ruler as a reference. The data was collected from three experiments’ lesion sizes of Fg-FEBs and PH-1 (Lei-Jie Jia et al, 2017). Significant differences in lesion length were consistently observed (**Fig 6A and B**). The brown lesions’ lengths were used as an index for coleoptile infection severity and, consequently, the pathogenicity of the isolates. The lesion sizes were significantly longer for the *Fg-S.maltophilia* isolate than for the PH-1 (**Fig 6A and** B), indicating a FEB caused increased pathogenicity.

**Figure 6.**
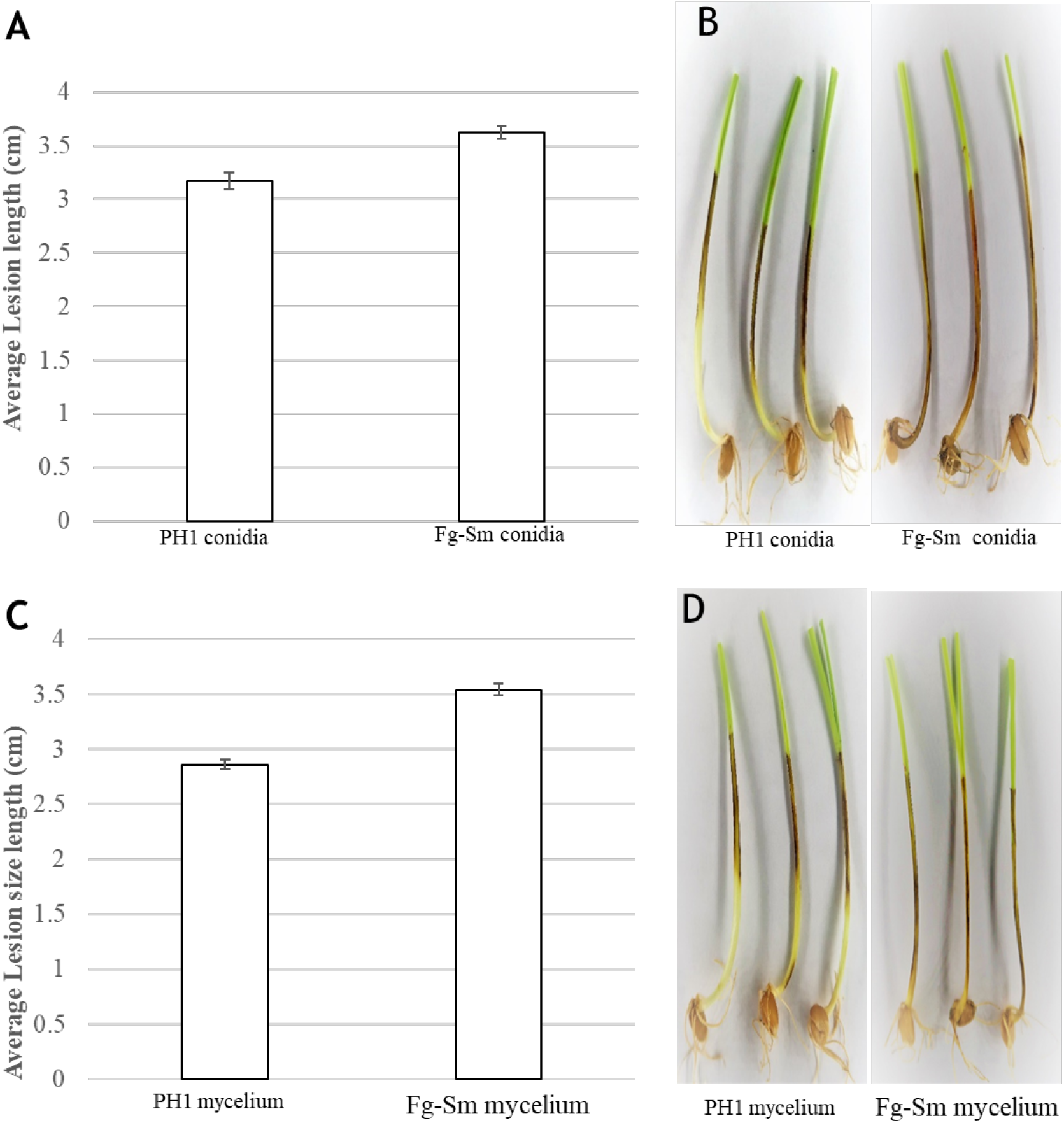
Pathogenicity assay of wheat coleoptiles infected with a conidia and mycelium of the Fg-*S.maltophilia* (Fg-Sm) isolate and PH-1. (A) Lesion length measurements of coleoptiles infected with conidia (B) Coleoptiles with lesions infected with a conidial suspension of PH1 and Fg-Sm isolate (C) Lesions length measurements of coleoptiles infected with mycelium. (D) Coleoptiles with lesions infected with mycelium of PH1 and Fg-Sm isolate. Error bars show 95% confidence intervals. Non-overlapping confidence interval bars are significantly different (P_same_≪0.05).

### 3. 7 Estimation of FEB load in F-S.*maltophilia* in culture conidia and infected plants

DNA was extracted from the wheat coleoptiles infected with conidia or mycelium of Fg-*S.maltophilia*. qPCR was performed to calculate the 16sRNA gene copies per fungal *His1mCherry*.Approximately 10 16sRNA gene copies per fungal *His1mCherry* genes were found in culture and coleoptiles when the plants were infected using mycelium as inoculum and around 6 when conidia were used. The main reason to use this qPCR comparison was to avoid detecting negligible amounts of bacterial genomes compared to fungal genomes. These differences were insignificant, although there were significantly more 16sRNA gene copies than mCherry copies in the biomasses in that the 95% error bars do not cross the 1/1 X-axis in Figure 7.

**Figure 7.**
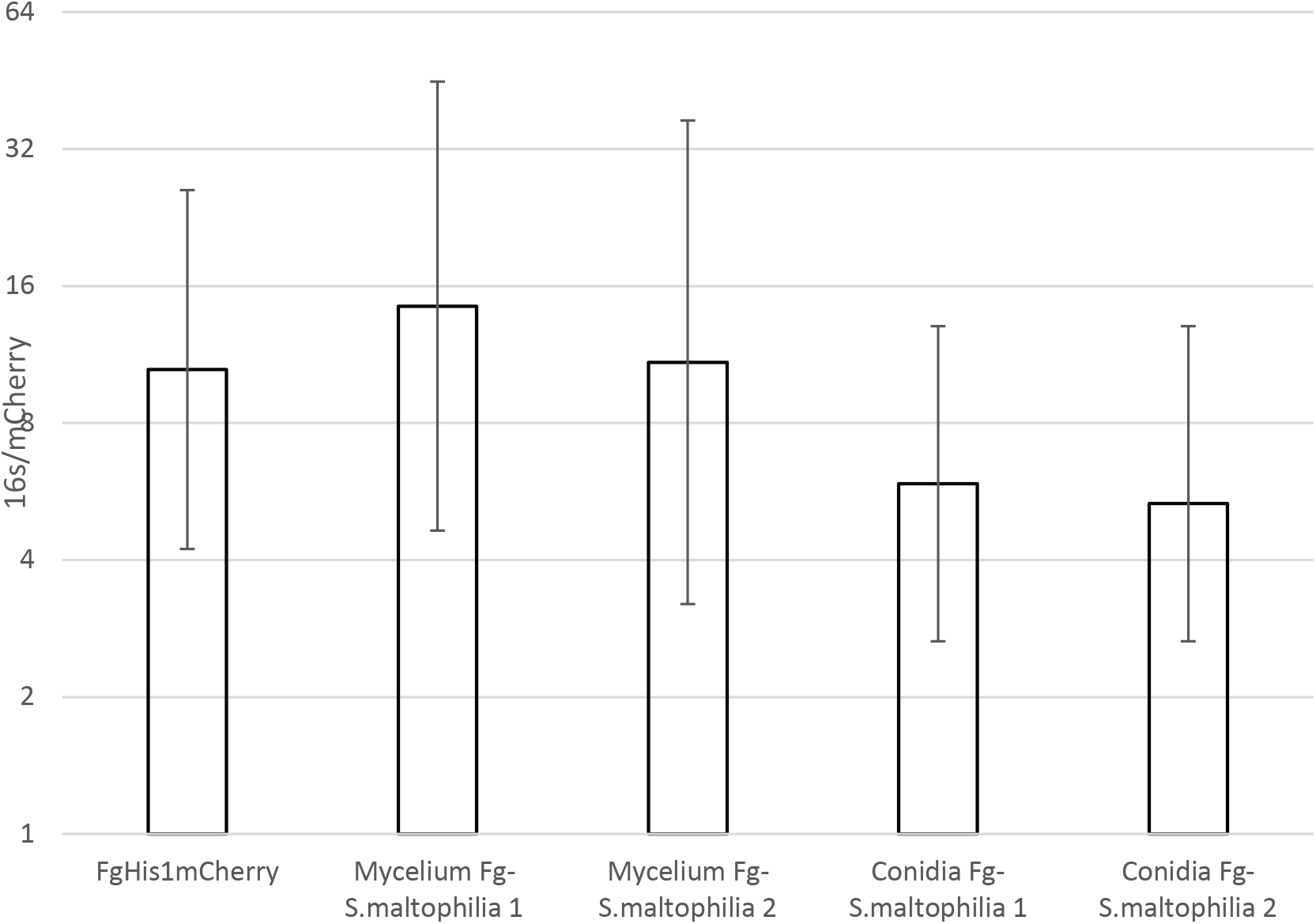
Estimation of *S.maltophilia* load in *Fg-S.maltophilia* during coleoptile infection. *In vitro* on Gentamicin (FgHis1mCherry) containing DFM and inside wheat coleoptiles infected with conidia or mycelium of *Fg-S.maltophilia* isolates and bacterial load estimated using qPCR. Each inoculation method was repeated two times independently with three replicates each time. Error bars show 95% confidence intervals. Non-overlapping confidence interval bars are significantly different (P_same_≪0.05).

### 3. 8 Nitrogen fixation capability of the *Fg-S.maltophilia* holobiont

We found several types of facultative Endohyphal bacteria, for example, *S.maltophilia, S. pavanii* and *P. ursingi*. All 3 are Gammaproteobacteria known as endophytes of plants and known for nitrogen fixation endophytes of plants and interestingly capable of nitrogen fixation (Ramos et al., 2011; Pillonetto et al., 2018; Alexander et al., 2019). The possible nitrogen fixation ability of the *Fg-S.maltophilia* holobiont was especially intriguing since that plant growth-promoting rhizobacterium (PGPR) species have recently been proven to increase peanut nitrogen fixation (Alexander et al., 2019) but also have plant growth-promoting effect on wheat under saline conditions (Singh and Jha, 2017). The potential nitrogen-fixing capability of *Fg-S.maltophilia* was tested by assessing the growth on a defined fungal medium without a nitrogen source added. Growth was tested on the nitrogen-free synthetic medium (modified DFM) with 2 times glucose concentration as carbon source, and no nitrogen source added. Gentamicin was added at a 512 μg/mL concentration to keep the FEBs inside the *Fg-S.maltophilia*.Dark red dense mycelium of *Fg-S.maltophilia* shows high biomass indicative of nitrogen fixation with a better yield on the medium single carbon source, glucose, and high formation of the secondary metabolite aurofusarin (Malz et al., 2005; Frandsen et al., 2006, 2011). In the dark red area of the *Fg-S.maltophilia*, the measured total extractable protein per colony area was for PH-1 3.5-10 times in the dense area than in a comparable area of PH-1, indicative of the nitrogen fixation (**Fig. 8**), while the protein content in the area outside the central dark red (**Fig. 8B**) was slightly lower than for PH-1 (**Fig. 8A**). The slower radial growth of the dark read area also indicates a higher biomass yield per glucose added made possible by the, possibly nitrogen fixing, carried bacteria. Very thin and fast radial growth similar to in PH-1 is seen for *Fg-S.maltophilia* outside the dark, red and dense centre, (**Fig. 8A and B**) most probably indicating growth of the fungus without N-fixation, similar to for PH-1. The red centre grows slower and later expands to cover the plate. Slower radial growth with dense mycelium generally indicates better yield on the available nutrients in the agar (Olsson, 1994, 1995, 2001).

**Figure 8.**
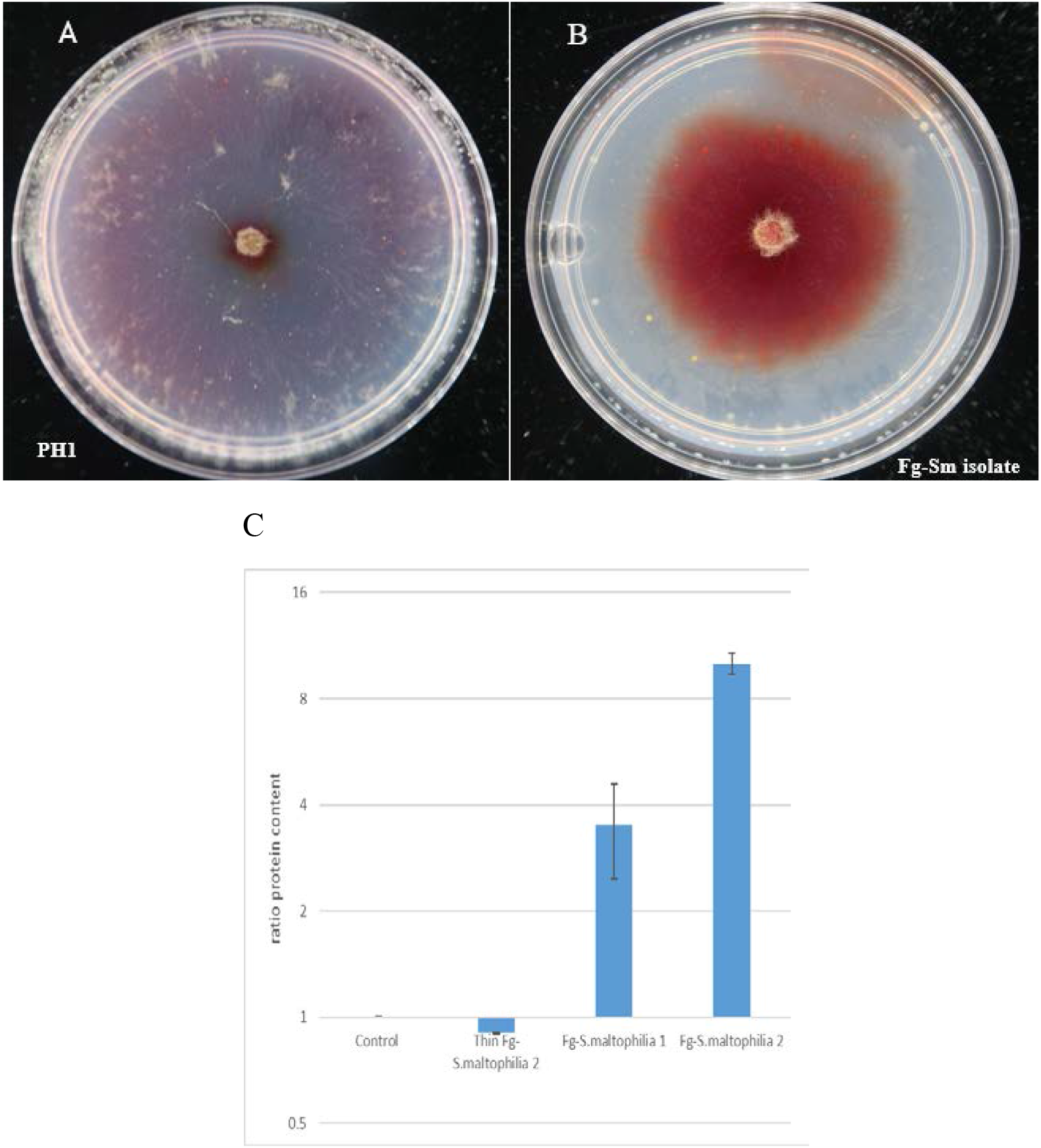
Nitrogen fixation capability of *Fg-S.maltophilia* v/s PH-1. Both were grown on a nitrogen-free synthetic medium supplemented with Gentamicin (512 μg/mL) with only glucose as a carbon source and no nitrogen source. Plates were extracted for total protein determination when the control cultures had grown to the plate edge. (A) At that time, the thin growth *Fg-S.maltophilia* cultures had reached the plate edge, as had the thin growth outer part of the *FgS.maltophilia* cultures. (B) The Image is from a few days later of plates that had not been used for protein extraction to measure the protein content, and confirmed that the dense red area expands over the thin area at the edge using the surplus carbon source that cannot be fully utilized without available nitrogen. (C) The ratio of protein content Fg-S.maltophilia (or PH-1 control)/PH-1 control in different areas. Two independent experiments were used to confirm the high protein content of the *FgS.maltophilia* deep red colony areas (Fg-S.maltophilia1 and 2). Error bars are 95% confidence intervals. Bars with non-overlapping error bars are significantly different (P(same)≪0.05).

## 4. Discussion

The soil bacteria we have found associated with *F. graminearum* as FEBs belong to Gammaproteobacteria, a bacterial class commonly found as facultative endohyphal bacteria of the fungal class Sordariomycetes that *F. graminearum* and many plant endophytes and plant pathogens belong to (Hoffman and Arnold, 2010). The FEBs load inside the fungus was stable over time when using gentamycin to kill bacteria escaping to grow on the medium (Baltrus et al., 2018). In our case, we calculated the bacterial load to avoid considering contaminating commensal negligible occurring bacteria as endohyphal. In future experiments, when we have sequenced the strains and know precisely how many gene 16sRNA gene copies there are in the bacterium, that can, combined with a Southern blot of the fungus for control of number of mCherry incorporations, be translated to bacteria/fungal nuclei. The copy number of 16sRNA varies between species (Větrovský and Baldrian, 2013) and with the growth strategy of the bacteria with higher copy number for faster possible growth rates in their natural environments (Klappenbach et al., 2000). The bacteria we found as FEB belongs to Gammaproteobacteria and are reported to have around 4 copies per genome (Větrovský and Baldrian, 2013). That should mean around 2.5 bacteria or more per fungal nuclei since the copy number of the mCherry gene in the fungal genome is at least 1. Since *F. graminearum* can be found in the soil plot at our university used for infection studies, it is probably a re-association of bacteria for PH-1 in our bacterial-free background fungal isolate, FgHis1mCherry derived from PH-1, we have detected.

Several of the bacteria we have found are known for their nitrogen-fixing ability. That applies to *S. maltophilia*, a known plant growth-promoting bacterium (Singh and Jha, 2017) that can improve the nitrogen status of peanut plants (Alexander et al., 2019). We could show that the holobiont Fg-*S. maltophilia* can grow well and form dense mycelium with high protein content on a defined agar medium without added regular nitrogen sources (**Fig. 8**). Nitrogen is a limiting factor for plant colonization by plant pathogenic fungi, including *F. graminearum* (Josefsen et al., 2012); thus, it is not surprising that the holobiont Fg-*S. maltophilia* showed more pathogenicity than the background *F. graminearum* strain FgHis1mCherry (**Fig. 6**).

Interestingly the load of Fg-*S.maltophilia* inside the fungal conidia and in the seedlings infected by mycelia were similar to what we find in culture (**Fig. 7**). Thus the FEB is likely, in this case, vertically transmitted, as has also been shown for several endohyphal bacteria of Zygomycota and Glomeromycota, some of which are obligatory endohyphal (Deveau et al., 2018). Consequently, we should expect to find many vertically transmitted FEB inside most fungi in nature if we look for them and use techniques for growing the isolates that stop the bacteria from escaping and competing with the fungus and appear as contaminants. Regular lab media (Oberhardt et al., 2015) are usually much richer, than the natural environments fungi and bacteria are adapted to live and grow in (Olsson, 2001), including the so-called rich rhizosphere of plants (Newman and Watson, 1977). When growing the *Fg-S.maltophilia* on distilled water agar only on nutrients available in the standard culture agar (not purified agar or agarose), the bacteria stayed inside the fungus, and no colonies could be seen on the agar, indicating that this is the case in nutrient-poor natural environments.

## 5. Conclusion

The natural state of *F. graminearum* is probably as an Fg-FEB holobiont. This holobiont has a changed phenotype and ecological niche compared to *F. graminearum* without a bacterial partner. Most fungi might, in nature, contain endohyphal bacteria affecting the physiology and ecology and the niche available for the fungal-bacterial holobiont. If so, we need to change our general practice for isolating, growing and studying many fungi in the lab to investigate better the nature and physiology of fungi in nature, including fungal pathogens.

## 6. Novelty of research

❖ FEBs are known for fungal endophytes but have not been studied for fungal pathogens.
❖ PH-1 in cultures is free from bacteria but acquires FEBs from natural soil environments, indicating that containing FEBs should be the expected growth state for *F. graminearum*.
❖ Both mycelia and conidial spores contain FEBs, and the FEB is stably associated with *F. graminearum*. Thus this indicates that the FEB-fungus holobiont can be a vertically spread entity subjected to natural selection as if it was a new strain.
❖ Containing a FEB increased *F. graminearum* pathogenicity on wheat.
❖ The FEB bacteria investigated belong to species known to fix nitrogen from the air.
❖ *F. graminearum* containing the *S. maltophilia* FEB grows well and makes high amounts of proteins compared to without FEB, with no nitrogen additions to the substrate, indicating that the Fg-*S. maltophilia* holobiont fixes nitrogen from the air that the fungus can utilize.

